# Functional Analyses of Histone Methyltransferases in Sea Lamprey Embryos Undergoing Programmed DNA Elimination

**DOI:** 10.64898/2026.01.08.698478

**Authors:** Kaan I. Eskut, Claire Scott, Cody Saraceno, Vladimir A. Timoshevskiy, Zachary D Root, Jeramiah J Smith

**Author notes:** Equal contribution (Kaan I. Eskut, Claire Scott, Cody Saraceno). Current address: Department of Pediatrics, Washington University School of Medicine, St. Louis, MO 63110. Current address: Department of Obstetrics and Gynecology, C.S. Mott Center for Human Growth and Development, Wayne State University School of Medicine, Detroit, MI 48202.

## Abstract

During early embryogenesis, the sea lamprey (*Petromyzon marinus*) undergoes a dramatic form of genome reprogramming wherein specific chromosomes are selectively eliminated from somatic progenitor cells. These programmatic elimination events effectively silence all genes on these chromosomes in all somatic cells. Previous studies in lamprey and other eliminating species have shown that epigenetic silencing marks are enriched on germline-specific chromosomes during programmed elimination. These silencing marks include the histone marks H3K9me3 and H4K20me3, which are respectively deposited by KMT1A/SUV39 and KMT5/SUV420 methyltransferases. To test whether lamprey homologs of these methyltransferases contribute to deposition of silencing marks on eliminated (micronucleated) chromatin and whether these marks contribute to the highly coordinated process of DNA elimination in sea lamprey, we used Cas9 gene editing, lightsheet imaging and RNA sequencing to investigate their potential roles in DNA elimination and more generally during early development. Analysis of knockout embryos for four histone methyltransferases show that these genes contribute to the deposition of repressive histone marks on elimination micronuclei, but are not essential for programmed DNA elimination *per se*. Analysis of later embryogenic stages suggests that these marks may contribute to interim silencing of germline-specific chromosomes, and reveals major impacts on post-blastula survival and development.

## INTRODUCTION

The sea lamprey reproducibly removes a portion of its genome from somatic cell lineages during development. This process, known as programmed DNA elimination (or more generally programmed genome rearrangement) has been observed in other lamprey species as well as other vertebrate (e.g. songbirds and hagfish) and invertebrate (e.g. roundworms and ciliates) taxa (Yao and Gorovsky 1974; Cleffmann 1980; Pigozzi and Solari 1998; Müller and Tobler 2000; Chalker and Yao 2001; Pigozzi and Solari 2005; Liu et al. 2007; Goday and Pigozzi 2010; Wang et al. 2017; Smith et al. 2021). In many of these species, including lamprey, programmed DNA elimination effectively prevents the somatic misexpression of a subset of genes that are thought to have evolved to function specifically in the germline (Smith et al. 2018; Smith et al. 2021; Timoshevskaya et al. 2023). Programmed DNA elimination can therefore be conceptualized as a form of epigenetic silencing. However, unlike other forms of epigenetic silencing, it is essentially irreversible during development.

While relatively little is known regarding the specific molecular mechanisms that drive programmed DNA elimination in vertebrates, it has become clear that DNA elimination often interacts with other modes of epigenetic silencing, including histone and DNA methylation. These interactions were first observed in the context of zebra finch spermatogenesis. In zebra finch, DNA elimination occurs both during embryogenesis and during spermatogenesis, where it has been studied in more detail. During spermatogenesis (but not oogenesis) the finch germline restricted chromosome (GRC) typically undergoes elimination. During spermatogenic elimination, the GRC is enriched for trimethylated H3K9 and H4K20 relative to the somatically retained chromosomes in addition to other transiently enriched markers (Schoenmakers et al. 2010; del Priore and Pigozzi 2014). It is, however, not yet known if these same histone methylation marks are associated with embryonic elimination in songbirds as these elimination events have only recently been observed (Dedukh et al. 2025).

In contrast to other vertebrates that are known to undergo programmed DNA elimination, lampreys produce large numbers of externally fertilized eggs. These externally developing embryos provide opportunities for manipulating early development and provide the ability to readily observe all stages of elimination. In sea lamprey, DNA elimination occurs between ∼36 and 60 hours of development and is nearly complete by the onset of gastrulation. In cells that are fated to become somatic tissues, GRCs are isolated from the somatically-retained chromosomes during anaphase and subsequently packaged into dense micronuclei. Paralleling observations of zebra finch spermatogenesis, sea lamprey elimination micronuclei were found to be strongly enriched for H3K9me3 (Timoshevskiy et al. 2016). This enrichment is followed by secondary enrichment of 5-Methylcytosine (5mC) and the subsequent degradation of DNA within micronuclei.

The methyltransferases responsible for deposition of H3K9me3 and H4K20me3 marks are conserved across eukaryotes. In mammals heterochromatic H3K9me3 is deposited by the genes *SUV39H1* (*KMT1A*), *SUV39H2* (*KMT1B*) and *SETDB1* (*KMT1E*) (Rea et al. 2000; Fritsch et al. 2010; Hyun et al. 2017), the latter of which is generally responsible for the deposition of H3K9me3 to silence endogenous retroviruses and transposable elements when complexed with sarcopterygii-specific KRAB zinc finger genes (Schultz et al.

2002; Matsui et al. 2010; Liu et al. 2014; Rosspopoff and Trono 2024), in addition to its role in methylating various non-histone targets (Fei et al. 2015; Guo et al. 2019; Wang et al. 2019; Rapone et al. 2023). In mammals, *KMT1A* and *KMT1B* act redundantly during embryogenesis, whereas *KMT1B* becomes restricted to testes later in development and mediates the silencing of sex chromosomes during the early stages of spermatogenesis (O’Carroll et al. 2000).

Two paralogous methyltransferases are also responsible for deposition of the H4K20me3 mark in mammals and many other vertebrates: *SUV420H2* (*KMT5C*) and *SUV420H1* (*KMT5B*) (O’Carroll et al. 2000). In mouse embryonic fibroblasts, *KMT5B* appears to be largely responsible for conversion of H4K20me1 to H4K20me2, whereas *KMT5C* primarily acts to convert H4K20me2 to H4K20me3 (Schotta et al. 2008). However, this simple relationship may not translate to tissues in general, as expression of *KMT5B* is more ubiquitous across mouse tissues, and knockout of *KMT5C* in mice did not completely ablate H4K20me3 from their tissues (Schotta et al. 2008). Additionally, the human homologs of *KMT5B* and *KMT5C* both seem to show a preference for H4K20me1 as a substrate *in vitro* (Wu et al. 2013) suggesting the possibility that they may act independently. A third KMT5 gene (*KMT5A*) is responsible for monomethylation of H4K20, which provides a necessary substrate for trimethylation by *KMT5B* and *KMT5C* (Wu et al. 2013; Southall et al. 2014; Weirich et al. 2016).

Because H3K9me3 and H4K20me3 are the first epigenetic marks known to be enriched on germline-specific DNA during programmed DNA elimination in sea lamprey, we hypothesized that these marks might be required for DNA elimination. To test this idea, we first identified a single lamprey ortholog of the two mammalian *KMT1A* and *KMT1B* paralogs, and three KMT5 homologs as candidates for emplacing H3K9me3 and H4K20me3 marks (respectively) on eliminated chromatin. Cas9 knockout of these proteins, in conjunction with immunohistochemistry, light sheet microscopy, and RNAseq analyses, was used to test for effects of knockout on the deposition of H3K9me3 and H4K20me3 marks and elimination of germline-specific DNA and more broadly resolve the roles of these marks during early embryogenesis.

## RESULTS

### Identification and phylogenetic analysis of lamprey KMT1A/B and KMT5 homologs

The lamprey genome contains one annotated homolog of the human *KMT1A/B* genes that we refer to hereafter as *KMT1*, one homolog of the *KMT5A* gene and two homologs of *KMT5B/C* genes (Smith et al. 2018; Timoshevskaya et al. 2023). To better understand the relationship between lamprey and gnathostome *KMT5B/C* homologs, we aligned *KMT5B*/*C* orthologs from several divergent vertebrate taxa and lancelet (used as an outgroup) and used these alignments to construct gene trees (**Figure 1**). A tree was also generated for KMT1 homologs to aid in resolving the relationships between a single agnathan KMT1 homolog and gnathostome paralogs.

**Figure 1.**
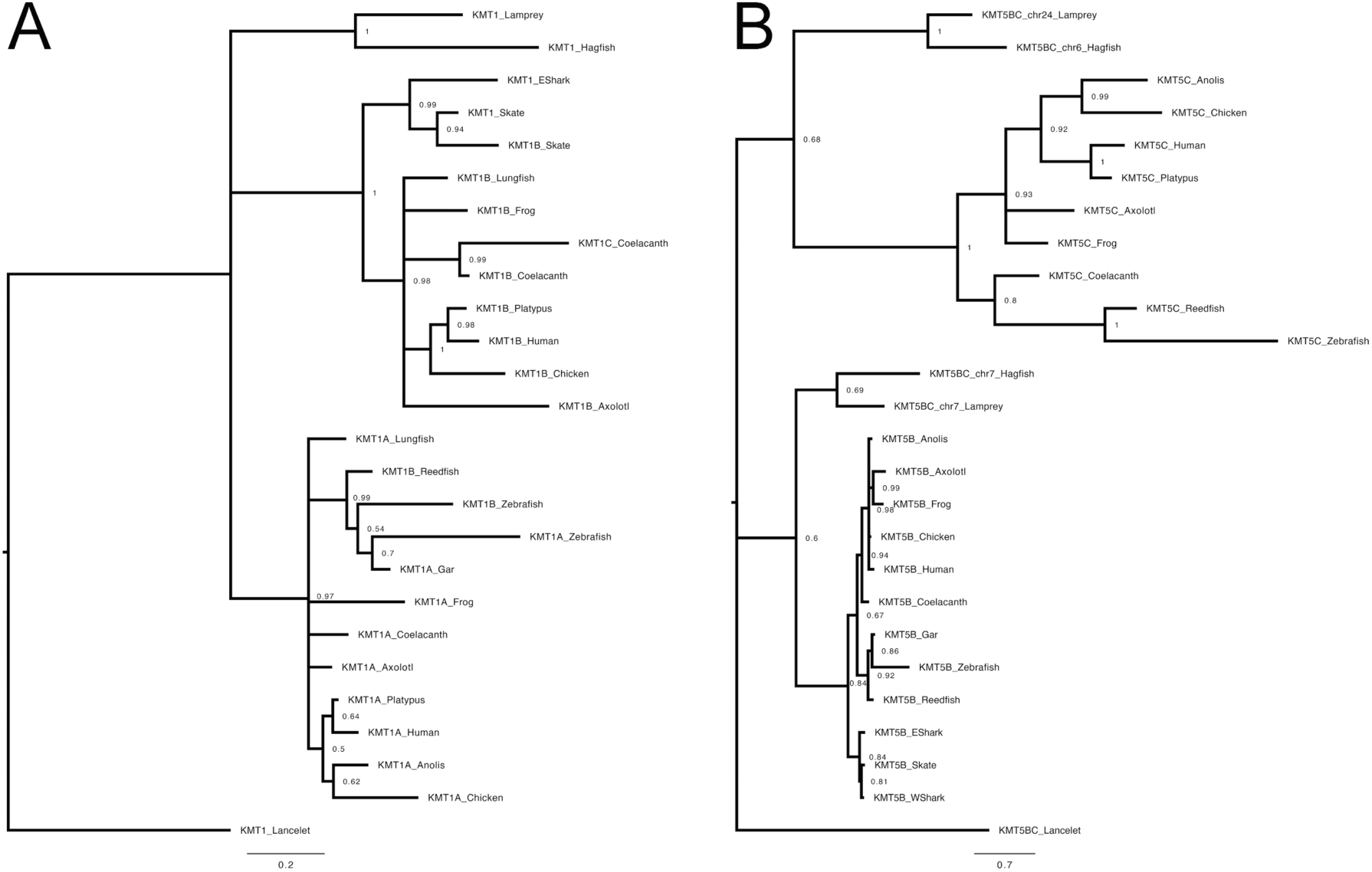
– A Gene tree illustrating the relationships between lamprey and gnathostome histone trimethyltransferase homologs. The evolutionary histories of chordate A) *KMT1* and B) *KMT5B/C* genes were inferred by using the PhyloBayes program under a CAT+GTR model, using Dayhoff 6-category amino acid recoding to account for potential bias caused by the high GC content in cyclostome genomes. The node labels are the Bayesian posterior probability scores. EShark = Elephant shark (*Callorhinchus milii*), WShark = Whale shark (*Rhincodon typus*).

In general, cyclostome homologs of *KMT5B* and *KMT5C* show relatively modest branch support for nodes connecting to gnathostome orthologs, with a homolog on lamprey chr7 (denoted *KMT5BC_chr7*) being more similar to gnathostome *KMT5B* orthologs and a homolog on lamprey chr6 (denoted *KMT5BC_chr24*) being more similar to gnathostome *KMT5C* orthologs. Low branch support values and topologies that are inconsistent with accepted vertebrate whole duplication events are frequently observed in trees that include data from cyclostomes, due in part to strong mutational biases over the long-term evolutionary history of the lamprey lineage (Smith et al. 2018; Marletaz et al. 2024). These biases have resulted in exceedingly high GC content of protein coding genes and constrain the mutational space that is explored by lamprey paralogs over evolutionary time, but are mediated to some degree by using Dayhoff6-recoding (Marletaz et al. 2024) (Figure 1, Figure S1). Notably, gnathostome *KMT5B* and *KMT5C* genes and their cyclostome homologs are present on paralogous chromosomes that are derived from the reconstructed ancestral chordate chromosome (CLGO) and were predicted to have diverged during the first vertebrate whole genome duplication (1Rv) based on phylogenetic signal from whole chromosomes (Marletaz et al. 2024). Overall, these trees reveal the impacts of duplication and paralog loss on the evolution of H3K9 and H4K20 tri-methyltransferases in the chordate lineage and confirm that agnathans possesses a single KMT1 ortholog and two KMT5B/C orthologs.

### Disruption of KMT homologs via Cas9 targeted mutagenesis

To further resolve the functions of KMT homologs, we disrupted the function of these genes in lamprey embryos using Cas9 to induce targeted mutations. The sole lamprey homolog of *KMT1* and three KMT5 genes were each targeted for knockout using two Cas9 guide RNAs that were positioned ∼50 bp apart within their 5’-most conserved exon, as assessed from multispecies conservation tracks displayed on the lamprey reference genome (Timoshevskaya et al. 2023), using the UCSC genome browser (Perez et al. 2025) and transcription/annotation tracks on SIMRbase (simrbase.stowers.org). For each gene, an initial set of five injected embryos and five of their uninjected siblings were fixed at 48 hours post fertilization (HPF) and imaged using light sheet microscopy to assess embryos for impacts of Cas9 injection on cell division rate or gross morphology. For the 10 control embryos used in this study, the average number of cells was 2278 (S.D. = 247.9, median = 2269), indicating that the average cell has undergone 11.1 cell cycles (**Figure 2A**). For the 20 crispant embryos, the average number of cells was 2414 (S.D. = 345.5, median = 2404), with each cell undergoing an average of 11.2 cell cycles. A comparison of cell number in crispant embryos revealed that, with the exception of *KMT5A*, all embryos fell within the developmental range of their respective controls (*KMT1* mean = 2410, S.D. = 351.5, Dunnett-test p-value = 0.16; *KMT5BC_chr7* mean = 2463, S.D. = 388.3, t-test p-value = 0.90; *KMT5BC_chr24* mean = 2166, S.D. = 320.5, p-value = 0.33). *KMT5A* embryos were at a slightly advanced stage of development compared to their control siblings (control mean = 2113, S.D. = 116.7; *KMT5A* mean = 2618, S.D. = 240.9; t-test p-value = 0.016), though this difference was less than a single cell cycle on average and was similar to KMT5BC sibling matched controls.

**Figure 2.**
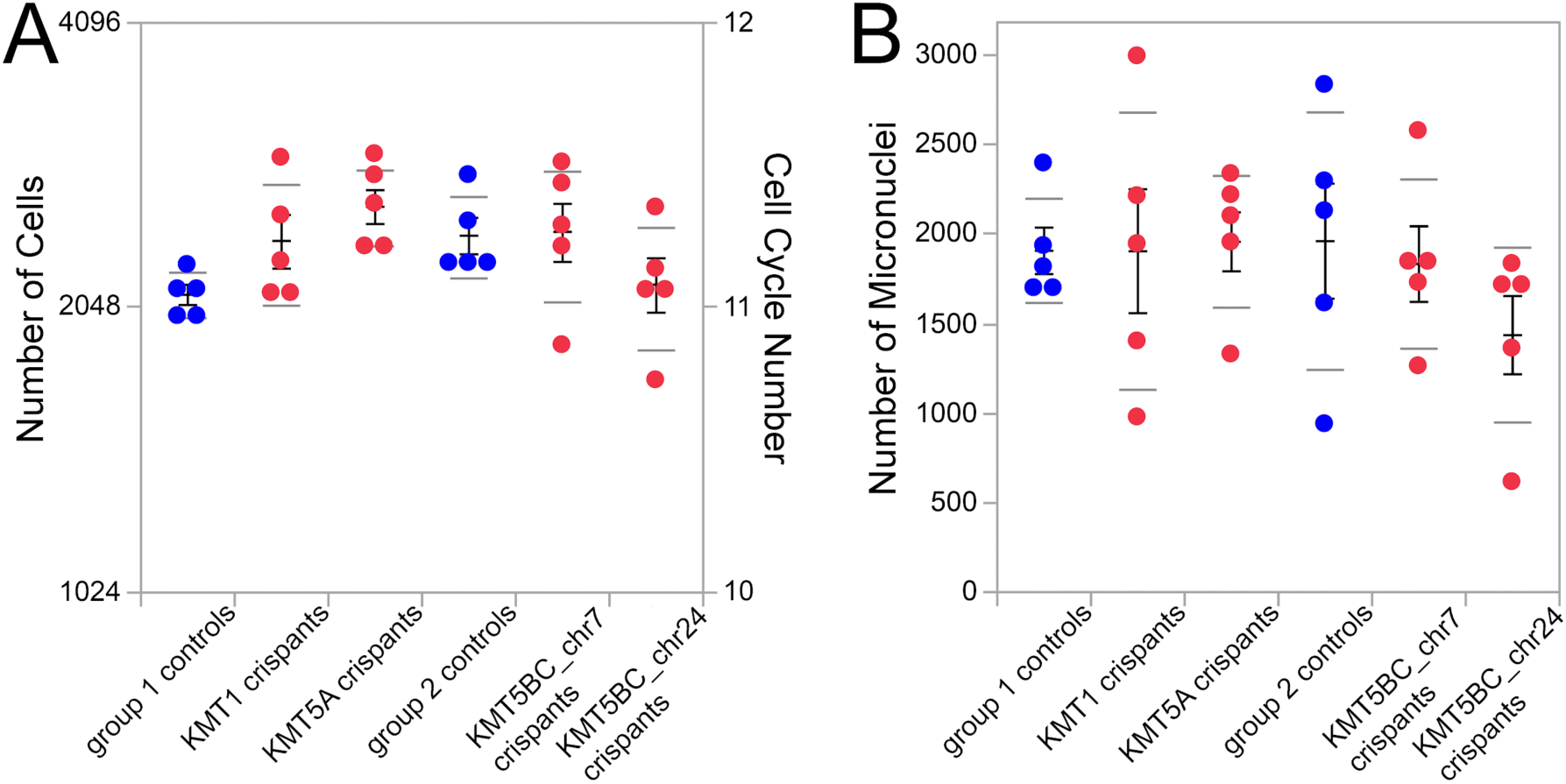
– Impacts of Cas9-mediated gene disruption on development and micronucleus formation in 48HPF lamprey embryos. **A)** Total number of cells present in each control and crispant embryo. Right axis shows the average number of cell cycles completed since fertilization. **B)** Total micronucleus counts do not differ significantly between SUV crispants and their respective controls. Standard error of the mean bars and standard deviation bars are shown in black and gray, respectively.

Following imaging, DNA was extracted from these same embryos and Cas target regions were amplified and sequenced to characterize the spectrum of Cas9-induced (and other) mutations. For each target gene, all five Cas9 injected embryos and two siblings were subject to DNA extraction, PCR amplification and Illumina sequencing, for a total of 28 embryos. Due to the challenges of extracting or amplifying DNA from individual fixed blastula-stage knockouts or the presence of large deletions, three individual embryos yielded only a small number of reads (and in one case none) but 24/28 embryos yielded more than 25 reads across the target region and most embryos were represented by several thousand reads (Figure S2).

Examination of mutation spectra from these reads revealed that uninjected sibling embryos were free of mutations in the target regions, although a few variants were detected outside of the target sites and these same mutations were observed to segregate among injected and uninjected siblings. Estimated indel frequencies were in excess of 90% for most injected embryos, indicating that Cas9 injection resulted in the efficient disruption of target sequences. Exceptions to this pattern were: 1) a nearly complete absence of mutations at one of two guide sites targeting *KMT5BC_chr7*, 2) an absence of observable mutations in several individuals with low read counts (noted above) perhaps due to a failure to sample large insertions or mutations encompassing the priming site or PCR failure, and 3) mutation frequencies of approximately 75% in two embryos (one *KMT5BC_chr24* injected and one *KMT5BC_chr7* injected). Indel frequencies ∼75% would be observed if one DNA molecule escaped editing following DNA replication at the first cell division.

Another notable feature of mutation spectra plots is the variability in the outcome of mutations induced by paired Cas9 guides. In this study, we used paired guide RNAs with the intent of inducing large deletions that resulted in the loss of the entire intervening coding sequence. Although four embryos (two KMT1 crispants and two KMT5A crispants) showed mutation spectra consistent with this intent, far more crispants were characterized by smaller indels that resulted from independent repair events. Despite the variability of repair outcomes in individual embryos, it appears that Cas9 injection into fertilized lamprey eggs provides an efficient means of inducing targeted mutations, consistent with other studies (Square et al. 2015; York et al. 2017; Yuan et al. 2018; York et al. 2021).

### Impact of Cas9 knockout on the formation of micronuclei

To test for potential impacts on programmed DNA elimination, we examined knockout and control embryos at 48HPF via lightsheet microscopy. At this stage, wildtype embryos consist of ∼2000 cells, with an average of 46% of cells containing one or more micronucleus. Nuclei and micronuclei were identified automatically using machine learning, and each micronucleus was assigned to a cell based on its proximity to the closest nucleus. For all 10 control embryos used in this study, the average number of micronuclei per embryo was 1937 (S.D. = 518.6, median = 1878). For the 20 crispant embryos, micronuclei were observed in all embryos at frequencies similar to their uninjected siblings, indicating that neither KMT1 deposited H3K9me3 nor KMT5 deposited H4K20me3 are required for DNA elimination. Among crispants, the average number of micronuclei per embryo was 1788 (S.D. = 544.6, median = 1772). Similarly, per-cell micronucleus counts and micronucleus volumes did not show any significant differences between crispants and their respective controls **(Figure S3)**.

### Resolving roles of lamprey KMT paralogs

Given the deep divergence time between lamprey and human and the presence of both gnathostome and agnathan duplicates, we sought to more directly assess the function of lamprey KMT1 and the three distinct KMT5 genes in depositing methylation marks. Although H4K20me3 had not been previously assayed in lamprey, fluorescent immunolabeling revealed that elimination micronuclei show strong enrichment for this mark in lamprey embryos in comparison to interphase nuclei (Figure 3, Figure S4). Mutant embryos and their wildtype siblings were screened for the presence of histone modifications (H3K9me3 for KMT1 and H4K20me1/2/3 for KMT5A/B/C) using embryos from the same injection series described above.

**Figure 3.**
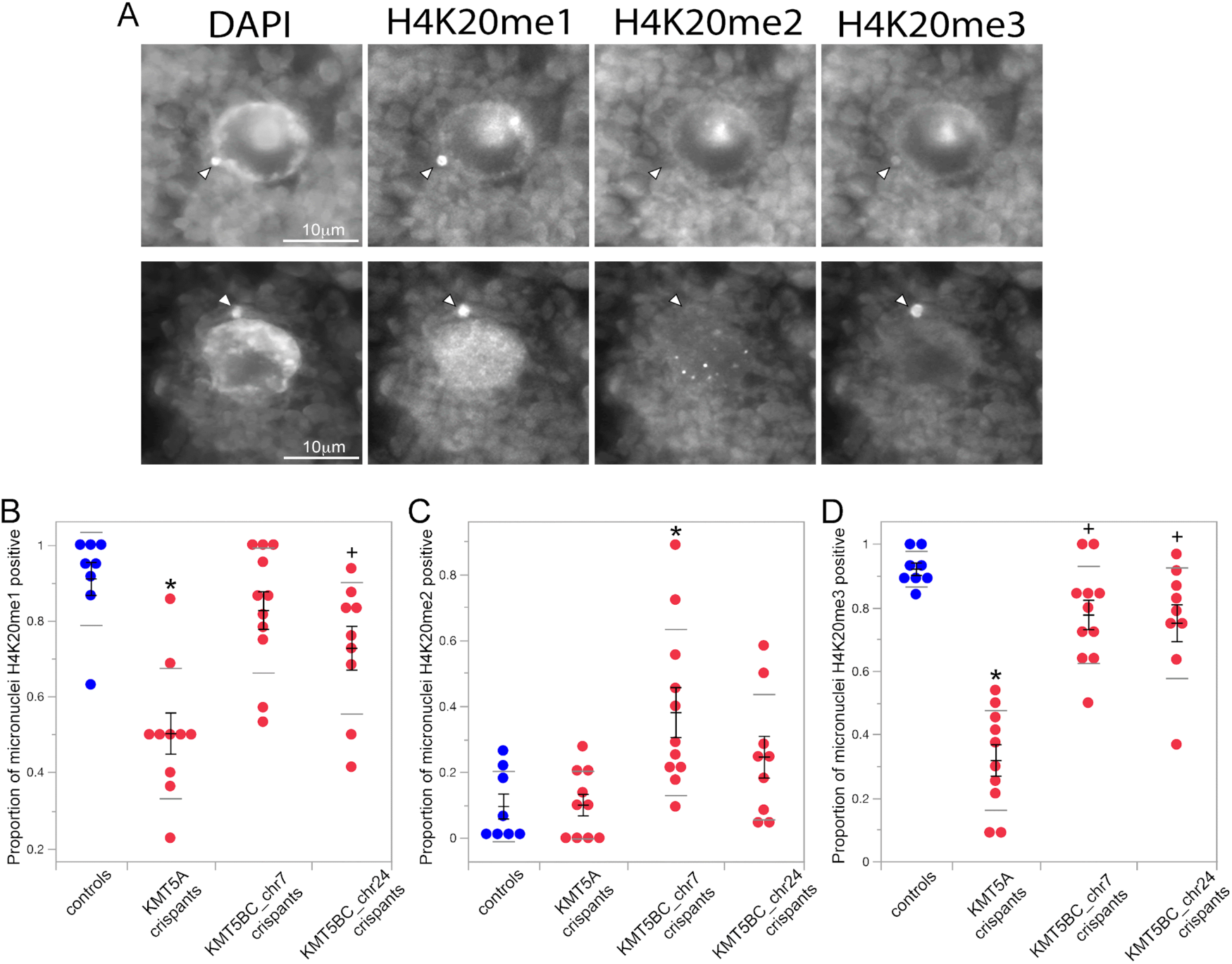
– Cas9 knockouts of KMT5 homologs alter the abundance of mono, di and trimethyl H4K20 on micronuclei. A) Examples of wildtype nuclei/micronuclei probed in H4K20 triple-labeling experiments. H4K20me1 and H4K20me3 are frequently enriched in micronuclei (arrows), whereas H4K20me2 is rarely observed in wildtype micronuclei (note the presence of nuclear puncta). B-D) Enrichment of H4K20me1,2,3 marks on micronucleated chromatin in wildtype and crispant embryos. Dots indicate the proportion of nuclei that are labeled for each mark for sections from individual embryos. Standard error of the mean bars and standard deviation bars are shown in black and gray, respectively. Symbols indicate groups that differ significantly from control siblings at: * = Dunnett’s t-test p<0.005, or + = p <0.1, >0.05.

**Figure 4.**
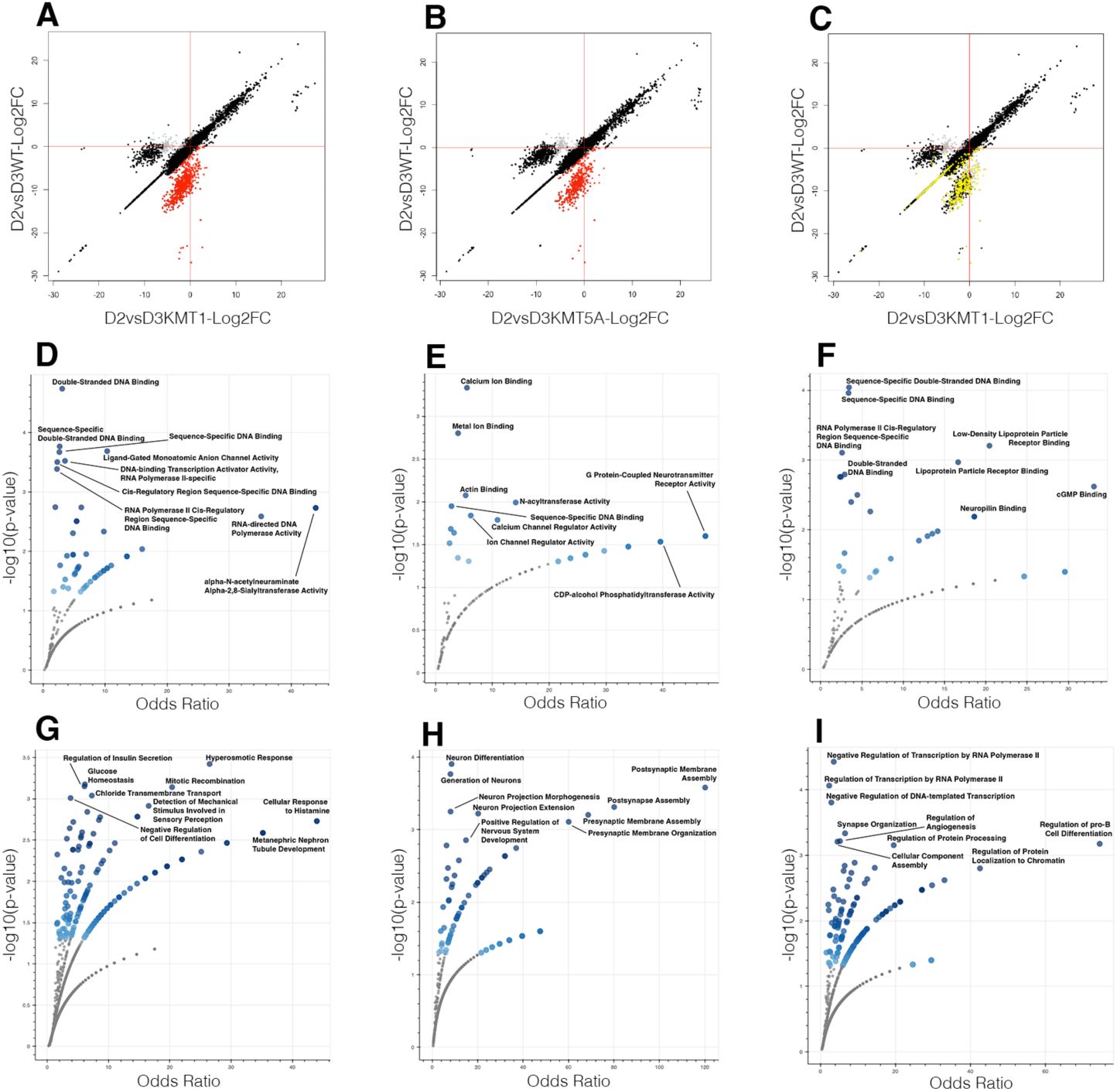
– Identification of unsilenced genes in *KMT5A* and *KMT1* crispants and gene ontology enrichment analysis of these gene sets. A, B, C) Comparison of gene expression changes in *KMT1* and *KMT5A* knockouts relative to wildtype. Each point represents a gene, with Log₂ fold changes (Log₂FCs) from the crispants are plotted on the x-axis and those from their wildtype siblings are plotted on the y-axis. Black points indicate genes showing statistically significant (p<0.05) differential expression, while grey points denote non-significant genes (p>0.05). Red points correspond to genes that are typically silenced in wildtype embryos but not silenced in crispants. Yellow points indicate the genes from *KMT5A* knockout unsilenced genes projected over the *KMT1* plot (highlighting shared genes). **D-I)** Volcano plots representing Gene Ontology (GO) enrichment analysis results. Each dot corresponds to an enriched GO term identified among the analyzed gene set. The x-axis denotes the odds ratio, reflecting the magnitude of enrichment, while the y-axis shows the-log10(p-value), representing the statistical significance of enrichment. The color intensity indicates the relative significance or enrichment score. This visualization highlights the most significantly overrepresented molecular functions and biological processes within the datasets. **D-F)** Enrichment of GO Molecular Function 2025 terms among unsilenced genes in: D) *KMT1* knockouts only, E) *KMT5A* knockouts only, **F)** both *KMT1* and *KMT5A* knockouts. **G-I)** Enrichment of GO Biological Process 2025 terms among unsilenced genes in: G) *KMT1* knockouts only, H) *KMT5A* knockouts only, **I)** both *KMT1* and *KMT5A* knockouts.

To test whether our candidate methyltransferases contribute to the deposition of H3K9me3 and H4K20me3 marks, we performed fluorescent immunolabeling of these marks on a separate group of sectioned Cas9 knockout embryos that were fixed at 48HPF. As was observed previously, H3K9me3 and H4K20me3 marks label a subset of micronuclei in 48HPF embryos (Timoshevskiy et al. 2016), whereas a smaller proportion of labeled micronuclei were observed in microinjected embryos. Cas9 knockout of *KMT1* resulted in a 46% reduction in the proportion of H3K9me3 labeled micronuclei (94/120 mni in controls vs 41/97 mni in crispants, χ^2^ = 16.1, p = 3.16^-5^), indicating that this KMT1 homolog plays a role in trimethylation of H3K9 in eliminated micronuclei. Disruption of all three KMT5 genes resulted in similar reductions in the number of H4K20me3 labeled micronuclei (75-76% reduction; Controls: 82/105 mni; *KMT5A*: 17/90 mni, χ^2^ = 40.4, p = 1.06^-10^; *KMT5BC_chr7*: 18/94 mni, χ^2^ = 41.8, p = 5.11^-11^; *KMT5BC_chr24*: 17/90 mni, χ^2^ = 40.4, p = 1.06^-10^), indicating that all three KMT5 genes contribute directly or indirectly to the deposition of H4K20me3. To further resolve the functions of KMT5 paralogs, we performed triple label IHC experiments on control and crispant embryos from each KMT5 target (**Figure 5**). As would be predicted on the basis of its sequence similarity to gnathostome KMT5A, knockout of the lamprey KMT5A ortholog results in a substantial reduction of H4K20me1 (controls: 193/208 mni, crispants: 87/166 mni, χ^2^ = 60.3; p = 4E^-15^): on average the proportion of immunolabeled micronuclei in crispants was 48.6 % that of their wildtype siblings.

**Figure 5.**
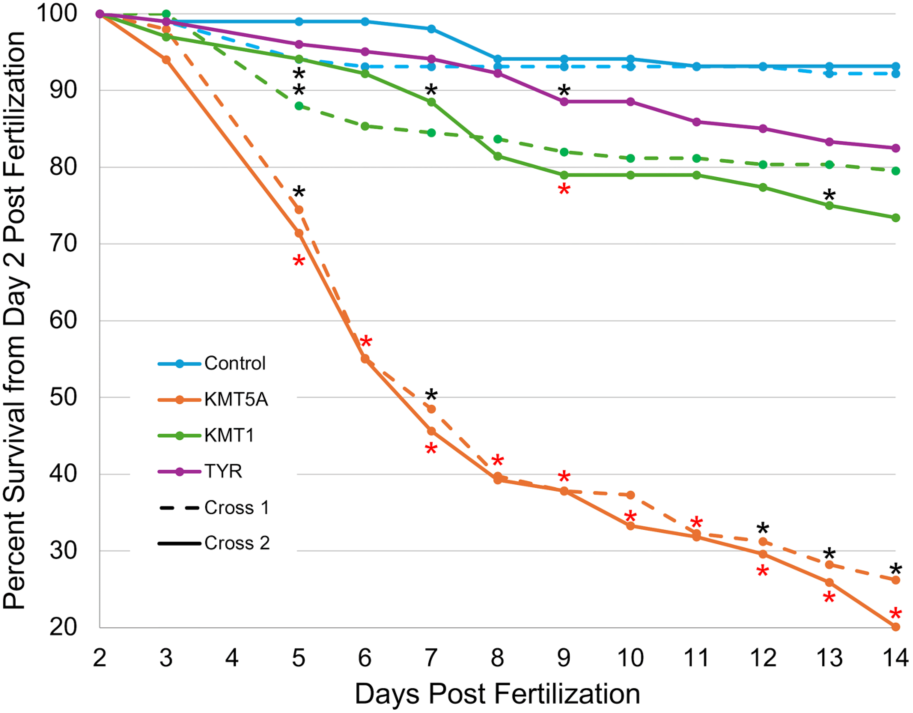
– Survival of KMT1 and KMT5A knockouts for the first two weeks post fertilization. Estimates of survival percentages are shown for knockout and control embryos for two separate crosses. Cross 2 includes an additional set of embryos that partially controls for generalized effects of Cas9 injection (using paired guides against the *tyrosinase* gene: *TYR*). Asterisks denote days where crispants had lower estimated daily survival rates in comparison to uninjected controls (Chi-square p<0.01). Red asterisks denote days where KMT crispants also had lower estimated daily survival rates in comparison to *TYR* crispants.

The other two KMT5 homologs appear to have somewhat overlapping roles in deposition of H4K20 methylation marks. Knockout of either *KMT5BC_chr7* or *KMT5BC_chr24* results in an increase in the proportion of H4K20me2 positive micronuclei, with this increase being slightly larger in *KMT5BC_chr7* crispants (controls: 19/208 mni, crispants: 71/211 mni, 3.68 fold; χ^2^ = 138.8; p = 2E^-32^) relative to *KMT5BC_chr24* crispants (crispants: 40/183 mni, 2.39 fold; χ^2^ = 32.4; p = 6E^-09^). In contrast H4K20me2 positive micronuclei were rare in control and *KMT5A* knockout embryos (20/166), consistent with the possibility that in this context H4K20me2 represents a transitional state between more stable monomethyl and trimethyl states in micronuclei. Outside of the micronuclei, H4K20me2 was observed to localize to puncta in wildtype interphase nuclei (Figure 6), consistent with its known role in double-stranded break repair (Paquin and Howlett 2018; Simonetta et al. 2018).

**Figure 6.**
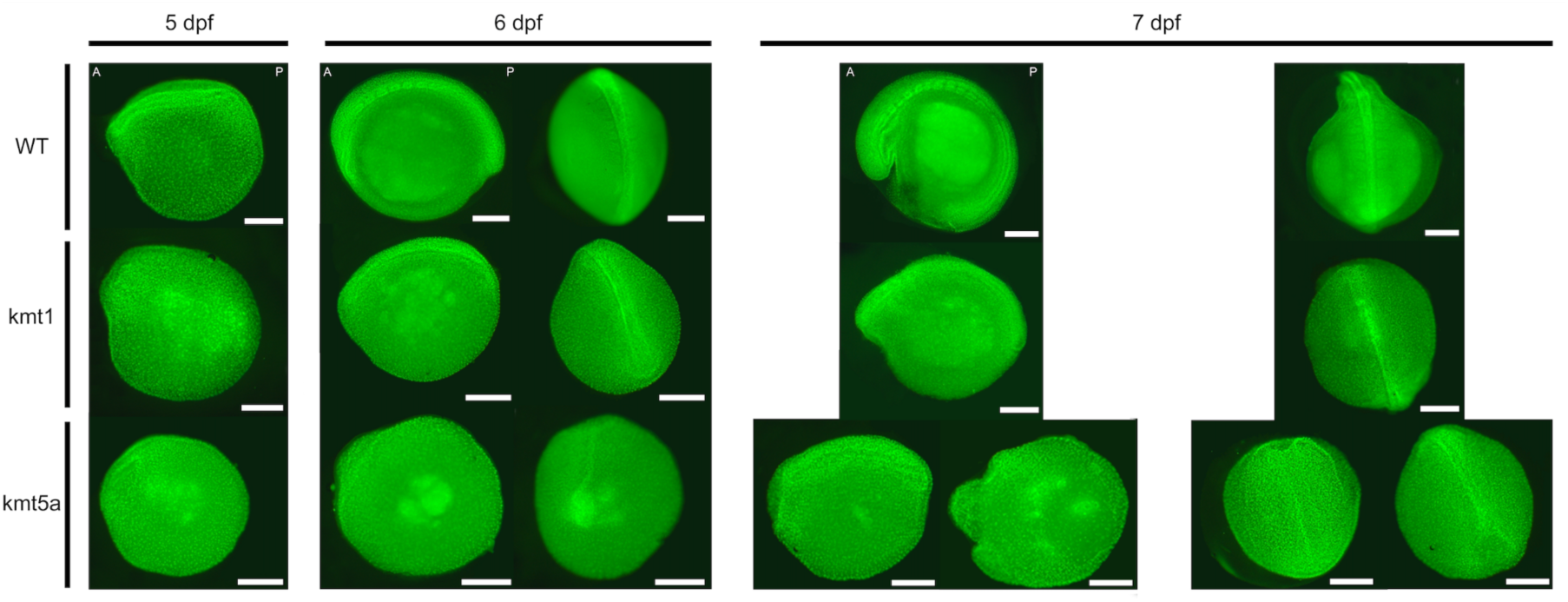
– Developmental impacts of *KMT1A* and *KMT5A* knockout during embryogenesis. Representative embryos at post-gastrula stages (5-7 DPF) are shown for wildtypes and two groups of crispants. Embryos at 6 and 7 DPF are shown in lateral (left) and dorsal (right) views. For *KMT5A* knockouts at 7DPF, two individuals are shown to showcase variation among crispants. *KMT1* knockouts show consistent developmental delays compared to WT, but more subtle morphological changes. *KMT5A* knockouts show more considerable developmental delays and more severe disruptions to morphology. Defects in neural tube/somite development were observed to varying degrees across *KMT5A* knockouts. Scale bars: 500 μm.

In triple labeling experiments only *KMT5A* knockout resulted in a substantial decrease in the proportion of H4K20me3 positive micronuclei, however it should be noted that the antibody used for these experiments differed from that used in our initial analyses (Figure 3) due to incompatibility of the initial H4K20me3 antibody with available antibodies for H4K20me2. The antibody used in triple label experiments has been reported to cross react with H4K20me2 (https://www.scbt.com/p/trimethyl-histone-h4-antibody-6f8-d9), which is consistent with the results observed here although H4K20me2 immunolabeling is very faint in wildtype micronuclei in comparison to H4K20me1 and me3 (Figure 3). In general, H4K20me3 immunolabeling was fainter in *KMT5BC_chr7* crispants relative to controls although the proportions of labeled nuclei were not significantly different (though still lower) when compared to controls (Figure 3). Taken together, histone IHC studies suggest that *KMT5BC_chr7* may have a somewhat greater impact on the transition from H4K20me2 to H4K20me3 relative to *KMT5BC_chr24*, reminiscent of the relationship between *KMT5B* and *KMT5C* mouse embryonic fibroblasts (Schotta et al. 2008).

### Impact of KMT Knockout on early zygotic gene expression

To understand the broader impacts of H4K20 and H3K9 methylation on early zygotic development, we also profiled gene expression in control and knockout embryos. Comparison of 3 days post fertilization (DPF) embryos to both age matched controls and 2DPF controls identified several genes that are silenced in controls at 3DPF (relative to 2DPF) but retain relatively high levels of expression in 3DPF crispants (Figure 4). We interpret this pattern of expression as indicating that these genes are normally silenced by H4K20me3 (for *KMT5A* crispants) or H3K9me3 (for *KMT1* crispants) during early embryogenesis. In total we identified 390 genes that appear to require H4K20me3 for silencing at 3DFP and 594 genes that appear to require H3K9me3, with 230 retaining 2DPF-like expression in both groups of 3DPF crispants (Supplementary Table 1). Notably, these unsilenced genes include several germline-specific genes (indicating incomplete silencing of germline specific chromosomes during later stages of elimination) and several transcription factors, including HOX genes, *POU3F3*/*OCT-8* and other transcription factors associated with nervous system development. Correspondingly, ontology enrichment analyses reveal that the targets of early silencing are associated with sequence-specific DNA binding, regulatory element binding, and neuronal functions (Figure 4, Supplementary Table 2-4).

### Impact of KMT Knockout on Embryonic Survival and Development

To better understand the impacts of KMT knockouts on later development, we reared embryos through the first two weeks of development and tracked daily survival. The survival of KMT5A knockout embryos was substantially lower than uninjected and controls injected with previously published gRNAs against *TYR* (Square et al. 2015). Surviving KMT knockout embryos were consistently delayed in development and showed a range of abnormal phenotypes (Figure 5,6). At 14 DFP, the remaining surviving KMT5A embryos had severe developmental defects, inconsistent with survival to free living stages of larval development. KMT1 crispants showed higher survival than their KMT5A crispant siblings, but lower survival than both their wildtype siblings and *TYR* crispant siblings.

## DISCUSSION

To better understand the relationship between DNA elimination and overlapping mechanisms of DNA silencing (H3K9 and H3K20 trimethylation), we identified and functionally characterized lamprey orthologs of mammalian *KMT1A/B*, *KMT5A* and *KMT5B/C* genes. Functional analyses show that these genes are responsible for establishing H3K9me3 and H3K20me3 marks on eliminating chromatin, but somewhat surprisingly, their loss appears to have little or no impact on the process of programmed DNA elimination. Expression analyses show that loss of these methylation pathways does impact the transient silencing of a small number of genes that are encoded on the GRCs, in addition to the silencing of several genes that are repressed by H3K9 and H3K20 trimethylation by 3DPF (early gastrulation). Analyses of later developmental stages show that both pathways have significant impacts on postblastula development and survival, consistent with studies in mouse.

### Evolution of KMT1 and KMT5 genes

The roles of KMT1 and KMT5 pathways in establishing methylation modifications on histones is highly conserved, despite differences in the retention of paralogous copies across major vertebrate clades. In mammals, H4K20 methylation in mammals is deposited by the *KMT5A* monomethyl transferase, and subsequently by *KMT5B*, and *KMT5C*. There appears to be overlap in the enzymatic activity of mammalian *KMT5B* and *KMT5C* with respect to deposition of H4K20me3 (Schotta et al. 2004; Schotta et al. 2008). Likewise, our analyses found that the two *KMT5B/C* homologs in lamprey also show functional overlap, with Cas9 disruption of either resulting in a partial decrease in H4K20me3 deposition and an enrichment in H4K20me2. The slightly stronger impacts of KMT5BC_chr7 with respect to H4K20 trimethylation is similar to that reported for mouse *KMT5B* in comparison to *KMT5C*. These functional similarities align with our gene phylogenies for chordate *KMT5B/C* homologs and suggest that the subtle functional divergence of *KMT5B* and *KMT5C* genes may have occurred early in the evolutionary history of the vertebrate lineage.

Perhaps not surprisingly, the history of duplication and retention of KMT1 homologs follows a pattern that differs from *KMT5B/C*. In contrast to mammalian genomes, which generally possess two divergent KMT1 orthologs, sea lamprey and hagfish genomes each possess a single identifiable KMT1 ortholog that is phylogenetically equidistant to mammalian *KMT1A* and *KMT1B* (Figure 1). Similarly, whole-chromosome phylogenies for the set of all derivatives of the ancestral chordate chromosome CLGE do not fully resolve the relationship between lamprey chromosomes and gnathostome chromosomes, but indicate that the chromosomes carrying gnathostome *KMT1A* and *KMT1B* genes likely duplicated prior to the split of the ancestral lineages leading to modern gnathostomes and agnathans (Marletaz et al. 2024). While phylogenetic analyses do not resolve lamprey KMT1 as orthologous to either *KMT1A* or *KMT1B*, these analyses do reveal key aspects of KMT1 paralog evolution in gnathostome lineages. Specifically, *KMT1B* orthologs appear to have been lost in the ancestral ray finned fish (actinopterygian) lineage, and conversely *KMT1A* appears to have been lost in the ancestral shark (chondrichthyan) lineages, with both *KMT1A* and *KMT1B* apparently arising in the ancestral vertebrate lineage but only persisting as retained duplicates in the sarcopterygian lineage (lobe finned fishes and limbed vertebrates). The independent retention of deeply diverged paralogs across vertebrate phylogeny mirrors the functional redundancy of *KMT1A* and *KMT1B* genes in mammals and presumably other sarcopterygian species.

### H3K9me3 and H4K20me3 in the context of programmed DNA loss

Our analyses confirm that H3K9me3 and H4K20me3 are enriched on micronuclei and that these marks are deposited by KMT1 and KMT5 genes. However, neither of these marks appears to be essential for programmed DNA loss as micronucleus counts, distributions, and size are not affected by the disruption of any of the KMT genes assayed in this study. This seems somewhat surprising given that the modifications established by these genes are the first modifications known to be enriched on micronuclei during programmed DNA elimination in sea lamprey (Timoshevskiy et al. 2016) and are also enriched in eliminated chromatin in other species (Schoenmakers et al. 2010; del Priore and Pigozzi 2014). A previous study of chromatin silencers in lamprey programmed DNA elimination showed that knockout polycomb repressive complex 2 genes altered micronucleus counts, distributions and size in sea lamprey, though not the ultimate elimination of DNA (Saraceno et al. 2024). By contrast, the only impacts on micronuclear phenotypes we detect in KMT knockouts is a reduction in their corresponding histone marks. We interpret these findings as indicating that the roles of H3K9me3 and H4K20me3 in programmed DNA elimination are likely relegated to parallel events that do not directly impact DNA elimination. Our RNAseq analyses provide some evidence that this may include silencing of gene expression from micronucleated chromatin, given evidence for incomplete silencing of a small number of GRC-encoded genes. Alternatively, our findings could also indicate the existence of more than one fully redundant pathway driving DNA loss, in which case knockout of a single pathway (e.g. H3K9me3-driven) would be fully compensated by another pathway.

### H3K9me3 and H4K20me3 in the context of early embryogenesis

In contrast to programmed DNA loss, analyses of gene expression and embryonic development reveal important roles of KMT1 and KMT5 pathways. RNAseq analysis of wildtype and crispant embryos reveals several genes that are normally silenced by these pathways early in development, and analyses of later stages reveal impacts of survival and development. Notably, there was substantial overlap in the genes that were found to be silenced by H3K9me3 (via KMT1) and H4K27me3 (via KMT5A). Considering the set of genes that are normally repressed by 3DPF but retain expression in knockouts, we found that 59% of the genes that were unrepressed in KMT5A crispants were also unrepressed in KMT1 crispants. Likewise, 39% of the genes that were unrepressed in KMT1 crispants were also unrepressed in KMT5A crispants, the lower percentage being due to the detection of a larger number of unsilenced genes in KMT1 knockouts. This pattern appears to be consistent with previous studies in mouse, which found that KMT1A/B-deposited H3K9me3 is required for deposition of H4K20me3 by KMT5B/C genes in mouse pericentromeric chromatin (Schotta et al. 2004).

Regarding development and survival through later stages of development, the most severe impacts were observed in knockouts of the KMT5A genes, which is responsible for depositing H4K20me1 marks that act as a substrate for KMT5B/C trimethyltransferases. As with RNAseq analysis, differences in phenotype between lamprey KMT1 and KMT5 knockouts roughly mirror previous studies performed in mouse. *KMT5B/C* double knockout mice (*SUV420H1*/*SUV420H2*) are born at submendelian rations, presumably due to embryonic lethality, and those surviving to birth die perinatally (Schotta et al. 2008). In contrast, mouse *KMT1A/B* double knockouts (*SUV39H1*/*SUV39H2*) experience somewhat lower rates of mortality at embryonic, perinatal and adult stages with approximately one third of animals surviving through adulthood, albeit with severe impairments to growth and development (Peters et al. 2001).

### Conclusions

From an evolutionary perspective, functional and phylogenetic analyses of lamprey KMT 1 and 5 genes show strong conservation of their corresponding histone trimethylation pathways in jawed vertebrates, despite multiple duplications and losses spanning several hundred million years of evolution and the evolution of other epigenetic pathways, including programmed DNA elimination. In addition to H3K9me3 and H4K20me3, a variety of other mechanisms of DNA silencing exist within eukaryote genomes, and these mechanisms frequently interact and depend on one another to control developmental processes. In species that undergo programmed DNA elimination (an extreme form of DNA silencing), colocalization of H3K9me3 and H4K20me3 with eliminating DNA have been observed in both lampreys and birds. Despite this correlation, we find that neither KMT1 nor KMT5-mediated trimethylation is essential for DNA elimination. These results, in conjunction with previously published analyses of PRC2 crispants, suggest that the mechanisms of programmed DNA loss are robust to perturbation of several individual major silencing pathways.

## METHODS

### Production of lamprey embryos and Cas9 injection

Eggs and sperm were collected from mature adults by stripping gametes into crystallization dishes containing 10% Holtfreter’s solution and incubated for 10 minutes to permit fertilization. After visually confirming activation, embryos were rinsed in distilled water to remove excess sperm and maintained in 10% Holtfreter’s solution at 18°C throughout development. Cas9 ribonucleotide complexes were generated by adding 5 μL 3μM gRNA (annealed crRNA and gRNA) for each guide sequence, 0.5 μL of 10μg/μL Cas9 protein and 1 μL of Dextran Fluorescein. Embryos were injected with ∼2-3 nL of this solution and screened for fluorescence at 24 hours post fertilization. For our four target genes, gRNA pairs were prepared by IDT with the following guide sequences (*KMT1*: ATCGCTTGTCGGAGATCTGT, GGCACAAATATATCTGAAAA; *KMT5A*: CAGGCGGCCAGACTCGGCTG, GGCAGTTACGCAAGTCCCGT; *KMT5BC_chr24*: GTACTGGTCGAGCACAAGCG, TCTTGTGCGTCTGGAAGCCC; *KMT5BC_chr7*: GTGAGTCTGGAAAGCCAGGT, ATCATTGTCGCAAAGCTCCT). At 48 hours post-fertilization, separate pools of knockout embryos and their uninjected siblings were collected into 2 mL centrifuge tubes and fixed in MEMFA Fixative for 1 hour rinsed in PBS, dehydrated in increasing concentrations of methanol, and stored in methanol at –20°C.

### Embryo clearing and staining

Prior to DNA staining, fixed embryos were embedded in hydrogel and cleared using a modified passive clarity protocol (Timoshevskiy et al., 2016). Briefly, embryos were gradually rehydrated in successive methanol/PBS dilutions (75%, 50%, 25% methanol), rinsed 3X in PBS and then placed in hydrogel monomer solution (5% acrylamide supplemented with 0.5% VA-044) at 4℃ overnight on a nutator. To allow for hydrogel polymerization, embryos were then incubated at 37℃ with gentle rotation for 3 hours. After incubation, the embryos were briefly washed with PBS and then changed into stripping solution (8% SDS in PBS) and incubated at 37℃ with gentle rotation for 5 days. Cleared embryos were then washed with PBS 5 times, changing solution each time and then transferred into staining solution (PBS, pH=7.4, 0.1 Triton X-100, 0.01% sodium azide). 2 μL SYTO 21 dye (5 mM in DMSO) per 1 mL staining solution was also added and the embryos were left protected from light at room temperature overnight, after which an additional 2 μL of SYTO 21 was added and incubated an additional 24 hours to ensure complete staining.

### Lightsheet imaging

For analyses of micronucleus counts and sizes, lightsheet imaging was completed for five embryos in each category: *KMT1* crispants, *KMT5A* crispants, *KMT5BC_chr24* crispants, *KMT5BC_chr7* crispants and two control groups. All embryos from each group were siblings from the same spawn, and the duplicate control groups were included to account for the potential for staining differences between control groups. After immersing embryos in RIMS solution for 30 minutes, 5 embryos were embedded in 2.5% agarose gel (made with prewarmed RIMS solution), pulled through a capillary tube and set at 4℃ for no less than 10 minutes. After successful embedding, the embryos were placed into the Zeiss Lightsheet Z.1 machine and extruded one at a time into the RIMS solution for imaging. Images were taken with a 5x objective lens with dual-sided laser illumination. Following imaging, the resulting image was split in half with the respective laser as the only source of illumination. These images included an extra 90 pixels on either side of the central x axis to account for alignment. These images were then stitched together after aligning in the x, y, and z planes. To obtain numerical results, pipelines to count nuclei, micronuclei, and annotate the associations between nuclei and micronuclei were run for each embryo through the Arivis software.

### Immunohistochemistry

MEMFA-fixed embryos were embedded in paraffin and sectioned. The sections were then washed in xylene three times and then in decreasing ethanol concentrations into PBS (100%, 70%, 50% ethanol in PBS). The slides were incubated at 37℃ overnight in sodium citrate. After a wash in PBS, blocking serum was applied for a 1-hour room temperature incubation covered with hybridization slips in a moist, dark chamber. The primary antibody 3% in PBS (H3K9 trimethyl polyclonal antibody A-4036 Epigentek, H4K20 trimethyl polyclonal antibody A-4048 Epigentek), H4K20 monomethyl polyclonal antibody A-4047 Epigentek, Anti-Monomethyl Histone H4 Antibody (5E10-D8) Alexa Fluor® 647, Anti-Trimethyl Histone H4 Antibody (6F8-D9) Alexa Fluor® 546) was then applied, covered with hybridization slips, and incubated at 4℃ overnight in the same chamber. Following this incubation, 2 washes in PBS and 2 washes in PBST for 10 minutes each before application of the 1/100 PBS dilute secondary antibody [Alexa Fluor-488 F(ab’)2 fragment of rabbit anti-mouse Ig G (H+L) (Thermo Fisher Sci A21204)]. The secondary antibody incubation was 1-2 hours at room temperature. The same washes of 10 minutes twice in PBS and twice in PBST were performed before the application of DAPI and subsequent imaging on Olympus cellSens Dimension software.

### DNA extraction and sequencing to confirm Cas9 target disruption

To assess for Cas9 disruption of target loci, DNA was extracted from all imaged embryos using the MagJET Genomic DNA Kit and subsequently used for PCR. PCR primers for each gene were designed using Primer3 (Untergasser et al. 2012) and are as follows: *KMT1*: F(ACTTGAGTGGACTGGAGTTGA), R(CCGGGTAATGTCAAGATCAGC); *KMT5BC_chr7*:F(AGGAGAACAGGGTGAA ATGG), R(AGGCCATGGTGTTTGAGAAC); *KMT5BC_chr24*: F(AGGTCACGCAAGAGAACCAG), R(AGGAGCTCTGCGAGACTGAC); *KMT5A*: F(GTAGCTTGGTCCTGCAGTTG), R(TGAGCTATCACATGTGGCCA). As much gel as possible was removed from each embryo before proceeding with the extraction kit. PCR was run with a 60℃ annealing temperature for 34 cycles with a final 30 second extension period after completion of cycling. The resulting amplicons were combined into seven separate pools (two pools of control embryos and five pools of Cas9-injected embryos), with one embryo from each Cas9 target being represented in each pool to permit full barcode deconvolution. Adapter/barcode ligation and sequencing was performed by GENEWIZ. Mutations in control and Cas9 injected embryos were assessed and summarized using CRISPResso 2.0.42 (Pinello et al. 2016).

### Phylogenetic tree construction

Protein sequences were downloaded from NCBI gene, or annotation databases for hagfish (Marletaz et al. 2024) and lamprey (Timoshevskaya et al. 2023), then aligned using Clustal W v2.1 (Larkin et al. 2007). The phylogenetic trees were constructed with PhyloBayes v4.1 (Lartillot and Philippe 2004; Lartillot and Philippe 2006; Lartillot et al. 2007) under a CAT+GTR model, employing Dayhoff 6-category amino acid recoding to mitigate compositional heterogeneity arising from the high GC content characteristic of cyclostome genomes. The resulting phylogenetic trees were visualized using figTree v1.4.4 (Rambaut 2010).

### Transcription and gene ontology enrichment analysis

Total RNA was extracted from two pools of approximately 100 snap frozen embryos for each condition, following the standard Trizol extraction protocol with the addition of 10 mM dithiothreitol (DTT) to the lysis reaction. Library preparation and sequencing were conducted by Novogene using their “Plant and Animal Eukaryotic mRNA” pipeline. These sequences and corresponding expression analyses were submitted to NCBI (GEO accession number GSE314474). RNA-seq reads from this study were aligned to the reference genome (GCF_048934315.1) using HISAT2 v2.2.1 (Kim et al. 2019). Gene expression estimates were obtained from these alignments using StringTie v2.2.3 (Pertea et al. 2015), and differential expression analysis was conducted using the DEseq2 R package v1.42.1 (Love et al. 2014). Gene set enrichment analyses were performed using the EnrichR web suite (Chen et al. 2013; Kuleshov et al. 2016; Xie et al. 2021) and its Appyters module with the genes that have identifiable human homologs for each set.

## COMPETING INTEREST STATEMENT

The authors declare no competing interests.

## ACKNOWLEDGEMENTS

This work was funded by grants from the National Institutes of Health (NIH) (R35GM130349) and National Science Foundation (NSF) (MCB1818012) to JJS. We acknowledge the support of the University of Kentucky High-Performance Computing complex, and the Oncogenomics Shared Resource of the University of Kentucky Markey Cancer Center which is supported by the National Institutes of Health (P30CA177558).

## AUTHOR CONTRIBUTIONS

Writing: KIE, CSa, JJS; data acquisition: KIE, CSc, CSa, ZDR, VAT; data analysis: KIE, CSc, CSa, VAT, JJS. conception of the study VAT, JJS.

